# Edge-Aware Graph Attention Networks for Interpreting Biophysical Mechanisms from Molecular Dynamics Simulations

**DOI:** 10.64898/2026.09.03.749137

**Authors:** Mohd Ahsan, Chinmai Pindi, Larry Rodriguez, Paaras Shah, Giulia Palermo

## Abstract

Graph attention networks (GATs) are emerging as powerful artificial intelligence (AI) tools for learning biomolecular dynamics, yet extracting mechanistic insight from learned attention remains challenging. Here, we present an interpretable AI approach that combines molecular dynamics prediction with mechanistic interpretation through attention-derived communication networks. We develop three edge-aware GAT models – Edge-Conditioned, Edge-Injected, and Edge-Gated – that differ in how edge information is incorporated during message passing. Relative to a standard GAT, our edge-aware models improve coordinate prediction while recovering complementary aspects of residue communication. Application to Chignolin and HIV-1 protease demonstrates their ability to characterize communication networks underlying protein folding, allostery, and mutation-induced functional remodeling. Analysis of the communication networks using graph-based descriptors of communication throughput, intensity, and relay revealed that the Edge-Conditioned model preferentially emphasizes communication hubs characterized by high communication throughput and intensity, the Edge-Injected model preferentially highlights high-intensity communication within functionally important regions, and the Edge-Gated model most clearly resolves long-range communication relay. Overall, this approach provides an interpretable AI strategy for uncovering the communication mechanisms that underlie folding, allostery, and long-range signal propagation in molecular machines, while also supporting AI-guided modulation of biomolecular function and rational protein engineering.

## Introduction

Protein function is governed not only by three-dimensional structure, but also by conformational dynamics and long-range communication between residues.^1,2^ Molecular dynamics (MD) simulations provide an atomistic view of these motions and can reveal how local perturbations propagate through a protein to regulate folding, binding, catalysis, and allostery.^2^ Yet extracting mechanistic insight from MD trajectories remains difficult. The relevant signals are often distributed across many residues, fluctuate over time, and involve transient contacts rather than persistent structural features. Classical approaches such as dynamic correlation analysis, Markov state models, and residue interaction networks have provided powerful tools for interpreting biomolecular dynamics, but they typically rely on predefined descriptors and may not fully capture nonlinear relationships encoded in the trajectory ensemble.^3,4^

Graph neural networks (GNNs) offer a natural framework for learning from biomolecular simulations because proteins can be represented as graphs, with residues or atoms as nodes and spatial contacts or physical interactions as edges^5–7^. Graph attention networks (GATs) are especially appealing because they assign different weights to neighboring nodes, offering a potential route not only for predicting molecular dynamics but also for identifying residue-residue interactions that underlie long-range communication and allosteric regulation.^8^ However, GAT models were originally developed for general graph-learning problems, where edges primarily define connectivity. In biomolecular dynamics, an edge is not simply an abstract link. It may encode distance, contact persistence, orientation, or non-covalent interaction strength, all of which shape residue-residue communication during biomolecular dynamics.

This distinction creates an important methodological challenge for interpretable machine learning in biomolecular simulations. When graph attention is used to infer residue communication networks, the physical meaning of the learned attention depends critically on how edge information enters message passing and attention weights^5,8^. If edge information is used only to define graph connectivity, the resulting attention maps may overemphasize transient spatial proximity or large-amplitude motion and may not fully resolve physically meaningful communication. This issue becomes critical when attention weights are used for interpretation. Consequently, the way edge features are incorporated into message passing becomes central not only to prediction accuracy but also to the interpretability of the resulting attention patterns.

Several GNN approaches have demonstrated the importance of incorporating edge or geometric information to improve molecular prediction^9–12^. Edge-conditioned convolutional networks use edge labels to generate filters, while molecular models such as SchNet, DimeNet, NequIP, and TorchMD-Net incorporate interatomic distances, directional information, or equivariant representations to improve prediction of molecular properties, energy and forces.^13–16^ In parallel, GATs have emerged as powerful message-passing models that learn adaptive attention weights between neighboring residues, offering an attractive approach for interpretable molecular learning. However, extracting biologically meaningful residue communication from learned attention remains challenging, as the relationship between attention and the molecular mechanisms encoded in conformational dynamics is not straightforward. This raises a fundamental methodological question: how should edge information be incorporated into graph attention to accurately capture biomolecular dynamics while yielding learned attention that reflects physically meaningful residue communication?

Here, we present an edge-aware GAT approach for evaluating the role of edge representations in interpretable graph learning of biomolecular dynamics (**Figure 1**). We develop and benchmark three graph-attention models – Edge-Conditioned, Edge-Injected and Edge-Gated – that improve coordinate prediction relative to a standard GATs while enabling mechanistic interpretation of learned molecular representations. These models differ in how edge information is incorporated during message passing. In the first, edge features contribute to the attention coefficients; in the second, they are incorporated directly into the transmitted messages; and in the third, they act as gates that regulate information transfer (**Figure 1c**). This design isolates the distinct roles of edge information in determining where the model attends, what information is transmitted, and how communication is regulated.

**Figure 1.**
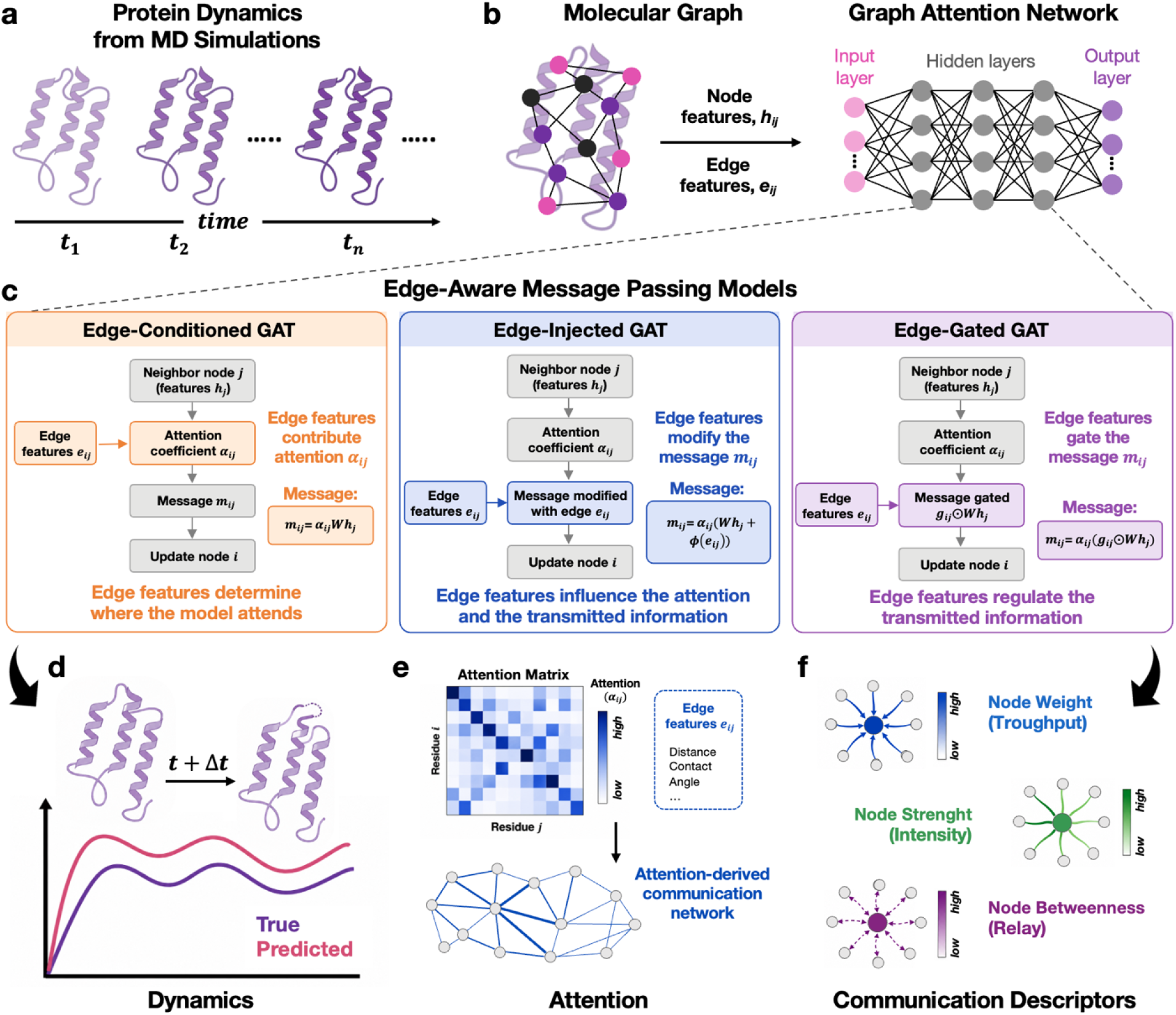
Edge-aware graph attention network for interpreting biophysical mechanisms from molecular dynamics simulations. **(a)** Molecular dynamics (MD) trajectories provide time-resolved protein conformations. **(b)** Each MD frame is represented as a molecular graph using residue-level node features ( ) and residue-residue edge features ( ), which serve as input to a graph attention network (GAT). **(c)** Three edge-aware message-passing schemes are introduced. In the Edge-Conditioned model (left), edge features contribute only to the attention coefficients ( ), determining where the network attends. In the Edge-Injected model (center), edge features are incorporated directly into the transmitted message ( ). In the Edge-Gated model (right), edge features regulate message propagation through an edge-dependent gate ( ). **(d)** The trained network predicts the protein conformation at the subsequent simulation frame ( ). **(e)** Learned attention coefficients ( ) are used to construct an attention-derived residue communication network. **(f)** Three residue-level communication descriptors are computed from these networks: (*i*) node weight, a measure of communication throughput, quantifying the cumulative attention-mediated communication flowing through a residue over the trajectory; (*ii*) strength, a measure of communication intensity, quantifying the magnitude of communication with neighboring residues during contact events; and (*iii*) betweenness, a measure of communication relay, quantifying the extent to which a residue functions as a bridge or bottleneck connecting communication pathways. Details in the text.

Because attention alone does not necessarily constitute a mechanistic explanation, we first evaluate the predictive relevance of learned attention using perturbation-based faithfulness analysis^17,18^. Validated attention weights are then transformed into time-dependent residue communication networks, which are characterized using three complementary graph-based descriptors – node weight, strength, and betweenness – to quantify communication throughput, local communication intensity, and communication relay, respectively. Through this approach, our edge-aware GAT models recover complementary aspects of residue communication. The Edge-Conditioned model preferentially identifies persistent communication hubs, the Edge-Injected model highlights high-intensity communication, and the Edge-Gated model most clearly resolves the long-range communication linking distant functional regions. Together, these components establish an interpretable AI framework for identifying biologically meaningful communication patterns from MD simulations.

We apply this approach to two systems of increasing complexity. Chignolin serves as a compact peptide benchmark for testing residue-level dynamics in a small folding system.^19^ HIV-1 protease provides a more complex allosteric benchmark, as its drug-resistance mechanism has been extensively characterized by prior MD studies and network analyses.^20^ Non-active-site mutations in HIV-1 protease are known to alter communication between distal mutation sites, the active site, and the flaps, producing indirect signaling pathways, weakened communication, and altered dynamics.^20^ Together, these benchmark systems allow us to evaluate not only the predictive performance of our edge-aware graph-attention models, but also how alternative edge representations shape mechanistic interpretation.

We show that learned attention can be transformed into biologically meaningful residue communication networks, providing an approach for uncovering molecular mechanisms from biomolecular dynamics simulations. This approach offers a general strategy for characterizing the communication networks that underlie biomolecular function – including folding, allostery, and long-range signal propagation in molecular machines – and for guiding the identification of residues governing these processes. More broadly, this approach is valuable for future uses of interpretable AI for modulating biomolecular function and rational protein engineering.

## Materials and Methods

Our approach integrates MD prediction and interpretability through three components. First, a graph-attention model was trained to predict residue-level structural evolution from MD trajectories. Second, the learned attention was evaluated by perturbation-based faithfulness analysis to determine whether it reflects features (e.g., residues or interactions) that are genuinely important for prediction. Third, attention weights were converted into dynamic residue networks and analyzed using three communication descriptors to characterize information flow. The overall workflow of our edge-aware graph-attention approach is summarized in **Figure 1**.

To enable a controlled comparison, all models shared the same graph representation, prediction task, training protocol, and evaluation pipeline. They differed only in how physical edge features are incorporated into the attention-based message-passing operation. This design allowed us to determine whether edge information simply biases attention, directly modifies the transmitted message, or regulates message transmission through a nonlinear gate.

### Graph-attention modeling of MD trajectories

#### Dynamic residue-level graph representation of MD frames

Each MD frame was represented as a dynamic residue-level graph that captures the instantaneous structural organization of the protein and its evolving residue-residue interactions (**Figure 1a,b**). The graph corresponding to simulation time *t* was defined as:

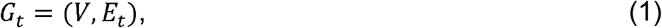

where *V* is the set of protein residues and *E_t_* is the set of residue-residue edges at frame *t*. Each residue was represented by its Cα atom. Node features, *h_i_*(*t*), encoded residue-level structural and dynamical information, while edge features, *e_ij_*(*t*), encoded the physical relationship between residue pairs. Because these quantities evolve throughout the trajectory, a new graph was constructed for every saved MD frame.

#### Edge-aware graph-attention models

The residue-level graphs were processed using GATs, which update node representations by aggregating attention-weighted messages from neighboring residues.^8^ For neighboring residues *i* and *j*, with node features *h_i_* and *h_j_* and edge feature *e_ij_*, the attention score was computed as:

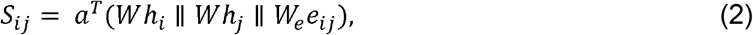

where *S_ij_* is the unnormalized attention score, *W* and *W_e_* are trainable transformations, *a* is a trainable attention vector, ∥ denotes concatenation. The corresponding normalized attention coefficient was calculated by applying a *softmax* over the neighborhood of residue *i*:

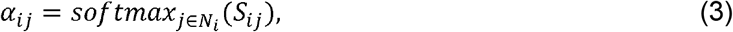

where *α_ij_* is the normalized attention coefficient that determines how strongly neighboring residue *j* contributes to the update of residue *i*, and *N_i_* denotes the neighborhood of residue *i*. The information transmitted from residue *j* to residue *i* was represented by a message *m_ij_*. The updated node representation was obtained by aggregating all incoming messages from the neighborhood of residue *i*:

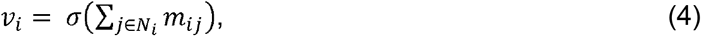

where *v_i_* is the updated representation of residue *i*, *m_ij_* is the message passed along edge *i*, *j*, and *σ* denotes a nonlinear activation function. The normalized attention coefficient *α_ij_* determines the relative importance assigned to neighboring residue *j*, whereas the message *m_ij_* specifies the information transmitted along edge (*i, j*). This distinction provides opportunities for incorporating edge information into the message-passing process.

Existing edge-conditioned and geometric molecular GNNs commonly use edge features to construct filters, encode interatomic distances and directions, or enforce equivariant message passing for molecular property, force, or potential prediction.^13–16,21^ Our objective was to isolate the role of edge features in attention-based message passing. To this end, we designed three graph-attention models that differ only in how edge features are incorporated into the construction of the message *m_ij_*. Specifically, edge information was used to: (*i*) bias attention coefficients, (*ii*) modify transmitted messages, or (*iii*) gate information transfer. All three models shared the same graph representation, network depth, prediction objective, and training protocol, allowing differences in predictive performance and learned communication patterns to be attributed solely to the role of edge information. The three edge-aware graph-attention models are described below and summarized in **Figure 1c**.

##### (i) Edge-conditioned attention model

(hereafter, Edge-Conditioned model, GAT-EC). Here, edge features contribute to the attention coefficient *α_ij_*, but not to the transmitted message:

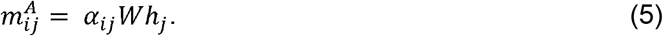

Thus, edge features determine where the model attends, while the transmitted information is derived exclusively from the neighboring node representation.

##### (ii) Edge-injected message passing

(hereafter, Edge-Injected model, GAT-EI). In this model, edge features are added directly to the transmitted message:

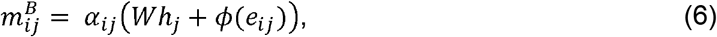

where *ϕ(e_ij_)* is a trainable transformation of the edge feature. In this design, edge features influence both the attention coefficient and the information transmitted between residues (i.e., the message).

##### (iii) Edge-gated message passing

(hereafter, Edge-gated model; GAT-EG). Here, edge features are used to gate the node-derived message:

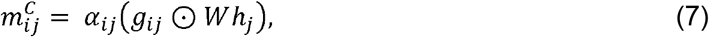

with:

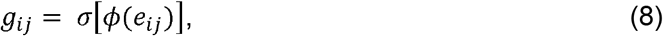

where *g_ij_* is an edge-dependent gate, σ is the *sigmoid* function, and ⊙ denotes element-wise multiplication. In this model, edge features act as a nonlinear filter that regulates information transfer between residues.

For all three methods, node representations were updated according to Eq. [4], followed by additional graph-attention layers and a final linear projection to predict the Cα coordinates.

#### Training objective and prediction evaluation

All models were trained using the same supervised one-step prediction task. For each MD frame *t*, the graph *G_t_* was used to predict the Cα coordinates of all residues at the subsequent saved frame, *t + Δt* (**Figure 1d**):

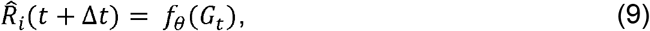

where *R̂_i_* (*t* + Δ*t*) ∊ ℝ^*N*x3^ is the predicted residue-coordinate matrix, and *f_θ_* denotes the trained graph-attention model. Thus, the model learns the short-time structural evolution of the trajectory by mapping each MD frame to its subsequent frame.

The model parameters were optimized by minimizing the mean-squared error (MSE) between the predicted and target Cα coordinates:

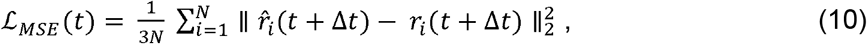

where *N* is the number of residues, *r̂_i_* (*t* + Δ*t*) is the predicted Cα coordinate of residue *i*, and *r_i_*(*t* + Δ*t*) is the corresponding coordinate from the MD trajectory.

After training, predictive performance was evaluated using RMSD-based analyses comparing the predicted and true Cα coordinate sets. We additionally compared the structural evolution of predicted and true trajectories relative to the initial MD frame to assess whether the model reproduced the overall trajectory-level dynamics. Before RMSD calculations, all structures were aligned by optimal rigid-body superposition.^22^ These analyses were used to verify that the trained model accurately captured the MD-derived structural dynamics before interpreting the learned attention weights were interpreted. Prediction accuracy therefore served as a prerequisite for interpretability. The mechanistic relevance of the resulting attention was then assessed independently using perturbation-based faithfulness and attention-derived residue networks.

### Validation of learned attention by perturbation analysis

Because neural attention does not necessarily provide faithful explanations, attention-derived interpretations should be validated independently.^17,18^ We therefore performed perturbation-based faithfulness analysis to test whether attention-ranked edges contribute more strongly to prediction than randomly selected edges.

For edge-faithfulness, edges were ranked according to their learned attention weights. A fraction *K* of the highest-ranked edges was masked by removing their contribution to message passing, and the prediction error of the perturbed model was recomputed. Masking was performed at *K* = 1, 2, 5, 10, 15, 20, 30, 40, 50 % of edges, and the prediction error was calculated for each mask. The increase in prediction error was defined as:

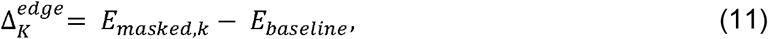

where *E_baseline_* is the prediction error of the unperturbed model and *E_masked,k_* is the error after masking the top-*K* attention-ranked edges. The same analysis was repeated with randomly selected edges as a control. The attention advantage was defined as:

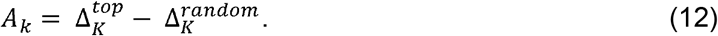

A positive *A_k_* indicates that masking high-attention edges degrades prediction more than masking randomly selected edges, supporting the predictive relevance of the learned attention ranking. *A_max_* denotes the maximum attention advantage observed across the evaluated masking percentages. To characterize the persistence of the attention advantage beyond its maximum (*A_max_*), we calculated *K*_50_, defined as the first post-peak masking percentage at which the attention advantage decreased to 50% of A_max_; when the crossing occurred between evaluated masking levels, *K*_50_ was estimated by linear interpolation. Smaller *K*_50_ values indicate that predictive information is concentrated in a smaller set of high-attention edges, whereas larger *K*_50_ indicate more distributed edge dependence. These perturbation tests quantify model dependence on attention-ranked edges; they do not, by themselves, establish biological causality.

### Mechanistic interpretation through attention-derived communication networks

Following faithfulness analysis, the trained attention weights were converted into time-dependent residue networks (**Figure 1e**). In these networks, nodes represent residues and edge weights represent attention-derived communication weights. Attention weights were extracted from the trained models, aggregated across attention heads and layers, and averaged over trajectory blocks. Because graph-attention weights are directional, they were converted into undirected residue networks before centrality analysis.

Three complementary network descriptors were used to characterize the learned communication patterns: (*i*) node weight, (*ii*) strength centrality, and (*iii*) betweenness centrality (**Figure 1f**). These metrices provide distinct and interpretable view of the learned residue communication.^23,24^

**(i)** The **node weight** of residue *i* was calculated directly from frame-level attention weights. For each frame, attention from all directed edges incident on residue *i* was accumulated and then averaged over the trajectory block (*T*):

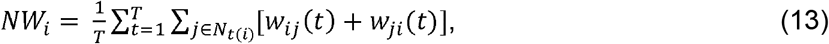

where *w_ij_(t)* is the attention-derived weight for the directed edge from residue *i* and *j* at frame *t*, and *N_t_(i)* is the set of residues connected to residue *i* in that frame. Edges absent from a given frame contribute zero attention weight. Node weight therefore quantifies the cumulative communication passing through a residue over the trajectory and is sensitive to both attention magnitude and contact persistence. Mechanistically, node weight quantifies how much attention-mediated communication flows through a residue over the course of the simulation and therefore serves as a measure of communication throughput.

**(ii) Strength centrality** was calculated from an undirected attention-weighted residue network. For each residue pair *(i, j)*, reciprocal directed attention weights were first averaged over the frames in which that edge was present:

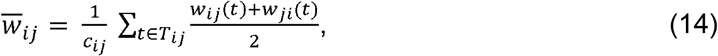

where *T_ij_* is the set of frames in which edge *(i, j)* is present, and *c_ij_ = | T_ij_|* in the corresponding edge count. The strength of residue *i* was then defined as:

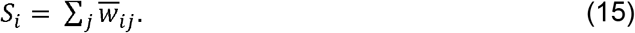

Strength therefore measures edge-level attention intensity, describing how strongly the model attended to edges associated with a residue when those edges were present. Unlike node weight, edge weights are normalized by edge occupancy, rather than by the full trajectory length. Consequently, node weight reflects how much communication is realized over time, whereas strength reflects how intense the communication is during contact events. Interpretation of node weight and strength therefore helps distinguishing persistent trajectory-level communication from intense couplings as they form. Mechanistically, strength quantifies how strongly a residue communicates with its neighboring residues when edges are formed and is therefore interpreted as a measure of communication intensity.

**(iii) Betweenness centrality** was used to identify residues that act as communication relays or bottlenecks within the learned attention network.^25^ Attention-derived edge distances were defined as:

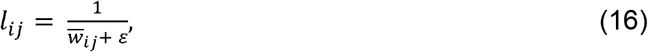

where ∊ is a small constant introduced to avoid division by zero. Betweenness was then calculated as:

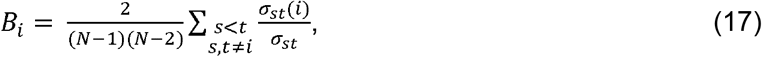

where σ*_st_* is the number of shortest paths between residues *s* and *t*, and σ*_st_(i)* is the number of those paths that pass through residue *i, N* is the number of residues. Betweenness therefore identifies residues that lie on dominant communication pathways and function as communication relays or bottlenecks. Mechanistically, betweenness thereby measures how much the network relies on a residue to relay communication between other^26–30^ residues and measures the capacity of a residue to act as a bridge or bottleneck. Together, these descriptors provide complementary mechanistic views of the learned communication network by quantifying communication throughput (node weight), communication intensity (strength), and communication relay (betweenness).

## Results

### Edge-aware GAT reproduce Chignolin dynamics and recover folding determinants

We first evaluated our Edge-Aware models on Chignolin, a fast-folding 10-residue β-hairpin peptide whose folding trajectory rapidly progresses from unfolded to folded states through intermediate misfolded states (**Figure 2a**). Its folding mechanism is well-characterized both computationally and experimentally, and provides a stringent benchmark for assessing both prediction accuracy and residue-level interpretability.^19,26–30^ Chignolin folding is stabilized by two structural determinants: the Tyr2-Trp9 cross-strand hydrophobic/aromatic interaction and a turn-region hydrogen-bonding network involving Asp3, Gly7, and Thr8.^19,31,32^ These features provide an ideal reference for evaluating whether attention-derived residue networks recover known physical determinants of folding rather than merely reproducing coordinate dynamics.

**Figure 2.**
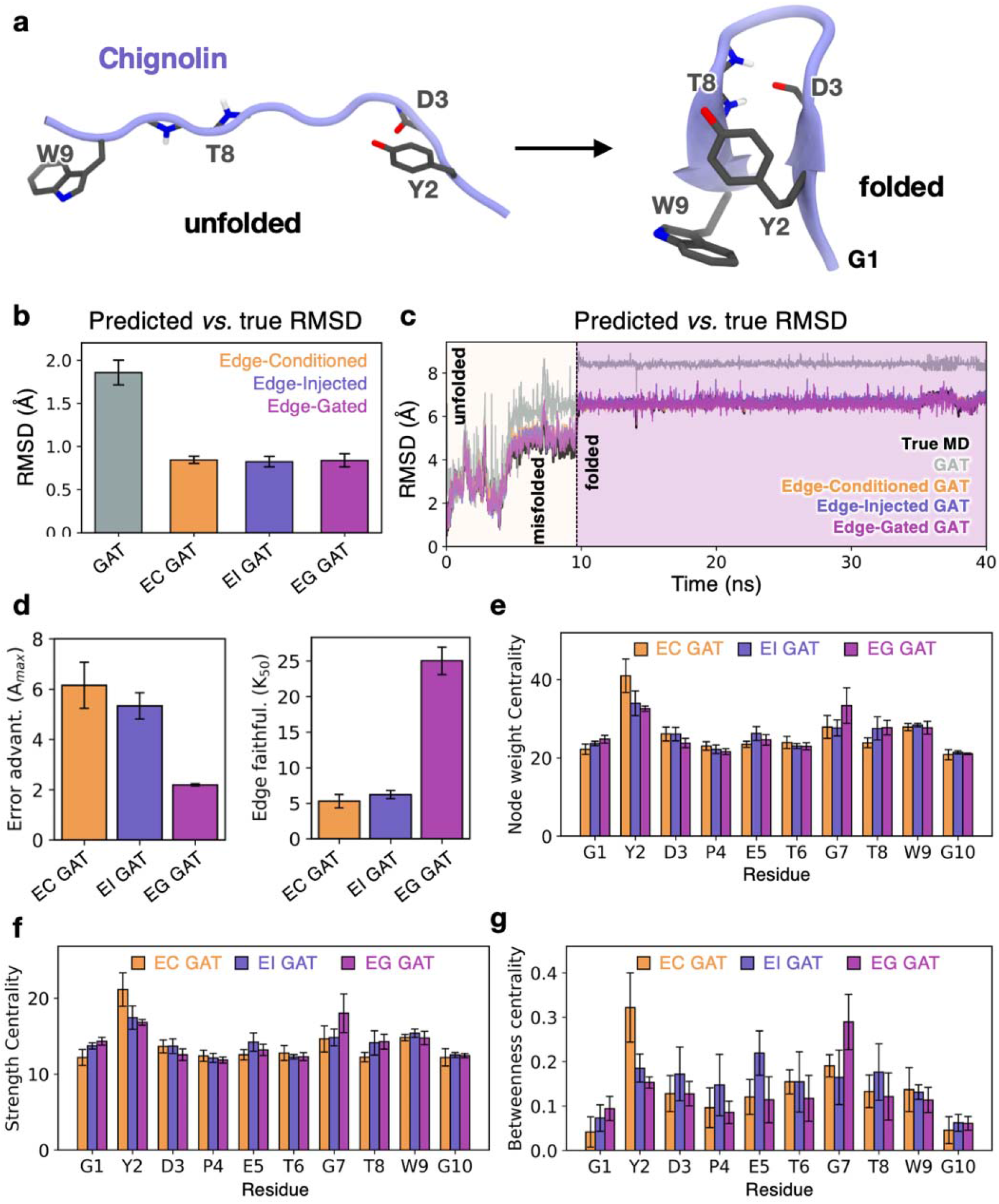
Edge-Aware graph-attention models reproduce Chignolin dynamics and recover residue-level folding determinants. **(a)** Overview of Chignolin highlighting the main structural determinants of the folded β-hairpin. The turn-region residues (Gly7, Asp3) and the terminal hydrophobic residues (Tyr2, Trp9) are shown. **(b)** Mean predicted–*vs.*–true Cα RMSD for the standard Graph Attention Network (GAT) and the Edge-Conditioned (EC GAT), Edge-Injected (EI GAT) and Edge-Gated (EG GAT) models. **(c)** Time evolution of the Cα RMSD relative to the first frame for the true MD trajectory and the corresponding trajectories predicted by the standard GAT and the three edge-aware models. The trajectory transitions from unfolded to folded conformations through intermediate misfolded states^26^. **(d)** Edge-faithfulness analysis for the three edge-aware models. The maximum attention advantage (*A_max_*, left) corresponds to the largest increase in prediction error obtained by masking high-attention edges relative to random edge masking, whereas *K*_50_ metric (right) is defined as the fraction of top-attention edges required to reach half of *A_max_* (details in Materials and Methods). Values in panels **(b)** and **(d)** are averaged over three independent simulation replicas. Error bars represent the standard error of the mean across replicas. **(e-g).** Residue-level attention-derived communication descriptors for Chignolin. Bars show the mean descriptor values for each residue using the three edge-aware models: **(e)** node weight, measuring communication throughput, **(f)** strength, quantifying local communication intensity, and **(g)** betweenness, measuring communication relay. Error bars in panels **(e-g)** represent the bootstrap standard error of the mean, estimated from 2,000 bootstrap samples generated using a two-level hierarchical block bootstrap procedure. Data for the Edge-Conditioned, Edge-Injected and Edge-Gated models are shown in yellow, blue, and magenta, respectively.

Prediction accuracy was evaluated using both predicted–*vs.*–true Cα RMSD and the RMSD relative to the initial frame. The standard GAT, which relies solely on graph connectivity without explicit edge-aware message passing, exhibited mean predicted–*vs.*–true Cα RMSD values of ∼1.4-1.5 Å, higher than the <1 Å achieved by all three edge-aware models (**Figure 2b**). The three models also accurately reproduced the time evolution of the Cα RMSD relative to the initial structure, closely tracking the conformational evolution observed in the MD simulations (**Figure 2c**). The standard GAT overestimated the RMSD throughout the trajectory, indicating that explicit edge-aware message passing is critical to capture the underlying collective conformational dynamics rather than simply propagating local coordinate changes.

We then performed perturbation-based edge-faithfulness analysis to test whether attention-ranked edges contributed more strongly to prediction than randomly selected edges. Specifically, we compared the prediction error resulting from masking high-attention edges with that obtained by masking randomly selected edges (see Materials and Methods). All three models exhibited positive maximum attention advantage (*A_max_*), supporting the predictive relevance of the learned attention ranking (**Figure 2d**). The Edge-Conditioned and Edge-Injected models exhibited larger *A_max_* values, indicating greater dependence on their highest-attention edges. Their correspondingly low *K*_50_ values (i.e., the fraction of high-attention edges required to reach half of *A_max_*, **Figure 2d**) indicate that this predictive information was concentrated within a relatively small subset of edges. The Edge-Gated model exhibited a smaller *A_max_* and a larger *K*_50_, indicating that predictive information was distributed across a broader set of high-attention edges. These results do not indicate superior or inferior edge faithfulness but rather demonstrate that the three edge-aware models encode predictive information through distinct communication strategies, ranging from a concentrated dependence on a smaller set of dominant edges (Edge-Conditioned and Edge-Injected models) to more distributed dependence (Edge-Gated model) across the residue interaction network.

Having established that the edge-aware models reproduced the MD trajectories and that their attention maps were prediction-relevant, we next examined whether the learned communication patterns recovered the known structural determinants of Chignolin folding. To this end, we characterized the learned communication network using node weight, strength, and betweenness (**Figure 4e-g**), which quantify communication throughput, communication intensity, and communication relay, respectively (details in Materials and Methods). The three models identified the principal folding motifs, while differing in how communication was distributed across the structural elements.

The Edge-Conditioned model organized its learned communication network primarily around Tyr2. Node weight (communication throughput), strength (intensity), and betweenness (relay) were all highest for Tyr2, while Trp9 and Gly7 also showed elevated values. These results recover contributions from both the Tyr2–Trp9 aromatic interaction and the turn region. The consistently dominant role of Tyr2 suggests that this model primarily organizes communication around the aromatic interaction.

The Edge-Injected model produced a more differentiated communication network. Communication throughput and intensity remained highest for Tyr2 while also highlighting Trp9 and Gly7, indicating strong communication through the Tyr2-Trp9 aromatic interaction and the Gly7 turn region. Relay shifted toward central and turn-associated residues (Glu5, Thr8, Asp3, and Gly7), indicating that these residues preferentially function as communication relays. This separation suggests that the Edge-Injected model distinguishes residues holding communication intensity from those that relay communication across the network.

The Edge-Gate model placed the Gly7 within the turn region at the center of the learned communication network. Communication throughput, intensity, and relay were all highest for Gly7, while Asp3 and Thr8 also contributed to the learned network, consistent with recovery of the Asp3–Gly7/Thr8 turn-stabilizing motif. Tyr2 and Trp9 remained prominent, particularly in local communication intensity, indicating preservation of the aromatic interaction, while this model prioritized the turn as the dominant organizing feature of the communication network.

Overall, the three models reproduced Chignolin dynamics and recovered the established folding determinants, while emphasizing different aspects of the communication landscape. The Edge-Conditioned model prioritized the Tyr2-Trp9 aromatic interaction, the Edge-Injected model distinguished communication hubs from relay residues, and the Edge-Gated model most clearly resolved the turn-stabilizing Asp3-Gly7-Thr8 motif as the dominant organizing center while retaining communication through the Tyr2-Trp9 interaction. Together, these results demonstrate that our edge-aware models recovered the established molecular determinants of Chignolin folding and provided a mechanistic interpretation of the underlying communication network.

### Edge-aware GAT reproduce HIV-1 protease dynamics and yield prediction-relevant attention maps

We next evaluated our edge-aware models on HIV-1 protease, a larger and more complex allosteric system. HIV-1 protease is a homodimer whose function depends on coordinated conformational dynamics, particularly flap opening and closing motions that regulate substrate access to the active site.^33,34^ Importantly, drug-resistance mutations can perturb these motions, despite occurring distal to the active site.^20,35^ We therefore compared the wild-type HIV-1 (WT HIV-1) protease with a non-active-site mutant (NAM, **Figure 3a**) derived from the multidrug-resistant clinical isolate CA84179, in which distal mutations have previously been shown to alter communication between mutation sites, the active site, and the flaps.^20^

**Figure 3.**
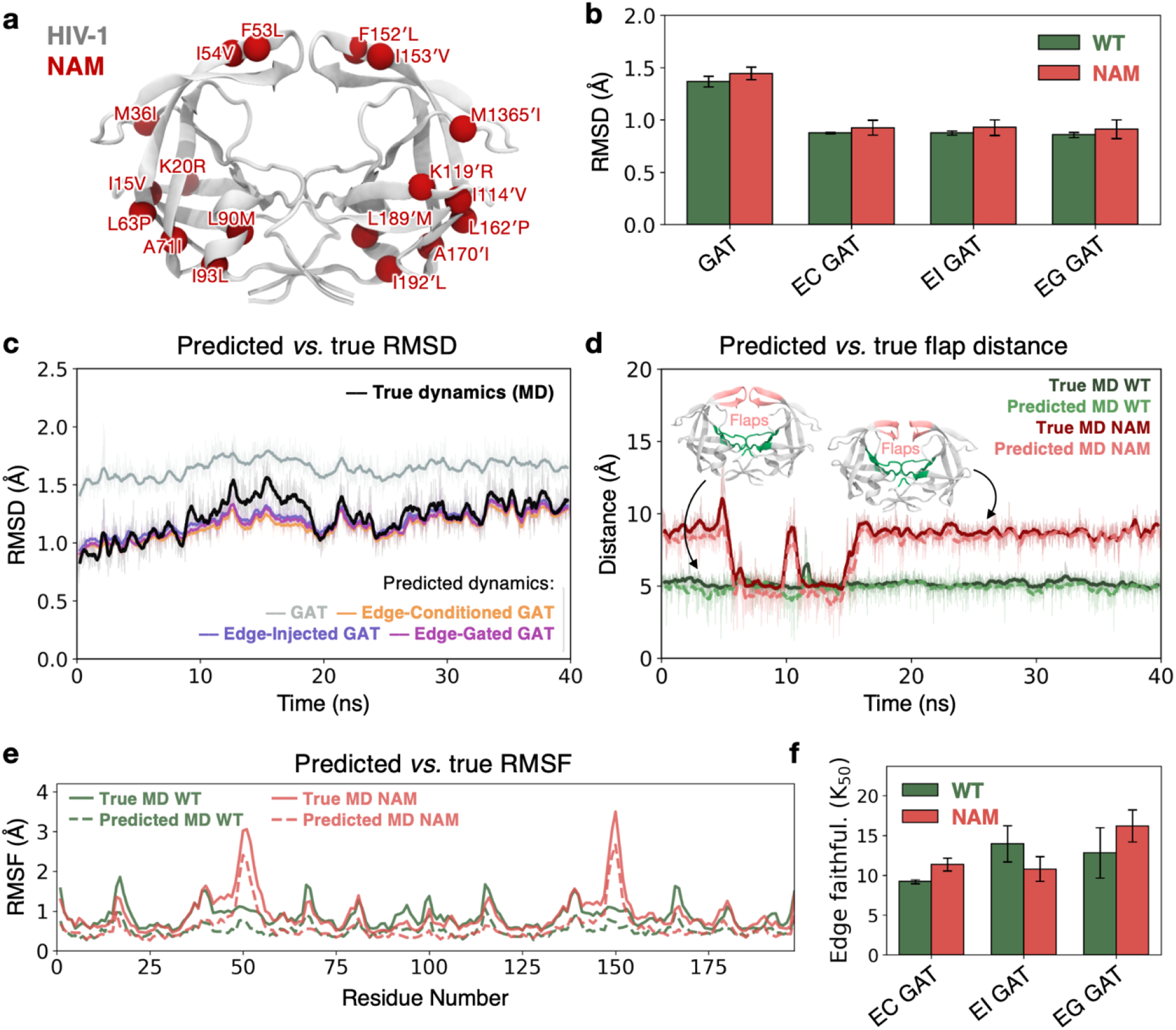
Edge-aware GAT models reproduce HIV-1 protease structural dynamics and generate prediction-relevant attention maps. **(a)** Structure of HIV-1 protease highlighting the mutations in the non-active-site mutant (NAM, red spheres). **(b)** Mean predicted–*vs.–*true Cα RMSD for the standard GAT and the Edge-Conditioned (EC GAT), Edge-Injected (EI GAT), and Edge-Gated (EG GAT) models for wild-type (WT) HIV-1 and NAM proteases. Error bars represent the standard error of the mean acros three replicas. **(c)** Time evolution of Cα RMSD relative to the first frame for the true MD trajectory and the corresponding trajectories predicted by the standard GAT and the three edge-aware architectures. **(d)** Time evolution of the flap tip-to-tip flap distance along the true MD simulation of the WT HIV-1 and NAM, and the corresponding trajectories predicted through the Edge-Gated model. In panels **(c-d)**, bold lines show 0.5 ns rolling averages and lighter traces the corresponding raw data. **(e)** Residue-wise Cα RMSF profiles for WT HIV-1 and NAM from the true MD simulations and Edge-Gated–predicted trajectories. **(f)** Edge-faithfulness analysis for the three edge-aware models. *K*_50_ is defined as the fraction of top-attention edges required to reach half-maximal attention advantage (*A_max_*) (see Materials and Methods). Values are averaged over three simulation replicas. Error bars are the standard error of the mean across replicas.

Prediction accuracy was assessed using complementary metrics of protein structure and dynamics. Analysis of the predicted–*vs.–*true Cα RMSD showed that, consistent with the Chignolin benchmark, the standard GAT model exhibited higher prediction error, with mean values of ∼1.4-1.5 Å, compared with values below ∼1 Å achieved by all three edge-aware models for both WT HIV-1 and NAM (**Figure 3b**). The three edge-aware models also accurately reproduced the time evolution of the Cα RMSD relative to the initial structure, closely tracking the conformational evolution observed in MD simulations (**Figure 3c**). In contrast, the standard GAT systematically overestimated the Cα RMSD. Beyond these global structural metrics, representative Edge-Gated model trajectories reproduced the flap tip-to-tip distance in both WT HIV-1 and NAM (**Figure 3d**), capturing the distinct conformational states and transitions observed in the MD trajectories. This indicates that the predicted dynamics preserve biologically relevant collective motions beyond residue-level coordinate prediction. Residue-wise RMSF profiles were also reproduced with mean absolute differences below ∼0.4 Å for both WT HIV-1 and NAM, indicating accurate prediction of residue-level fluctuations (**Figure 3e**). Together, these results demonstrate that our edge-aware models accurately capture both the global structural evolution and the functionally relevant conformational dynamics that distinguish WT HIV-1 and NAM.

We next performed perturbation-based edge-faithfulness analysis to validate the predictive relevance of the learned attention weights (details in Materials and Methods). The three edge-aware models exhibited positive maximum attention advantage *A_max_*, demonstrating that masking attention-ranked edges degraded prediction more than masking randomly selected edges. Edge-faithfulness analysis was further summarized using the *K*_50_ metric (**Figure 3f**). Across both WT HIV-1 and NAM, all three edge-aware models exhibited *K*_50_ values ranging from ∼9-16%, indicating that masking a fraction of high-attention edges had a substantial impact on prediction than masking an equivalent fraction of randomly selected edges. This validation shows that our explicit edge-aware message passing improves prediction of HIV-1 protease dynamics while generating prediction-relevant attention maps suitable for mechanistic interpretation. We next examined how the three edge-aware models organize residue communication within the WT HIV-1 and NAM mutant proteases.

### Residue-level attention changes identify mutation-sensitive sites in HIV-1 protease

We next examined how the non-active-site mutations remodel residue communication within the functional flap and active-site regions of HIV-1 protease (**Figure 4a**). Although the NAM substitutions occur distal to the catalytic center (**Figure 3a**), previous network analyses have shown that they perturb communication pathways linking the mutation sites, active site, and flaps.^20^ For each residue, we calculated *Δ*(W*T — NAM*), where negative values indicate enrichment in NAM and positive values indicate attenuation in NAM (i.e., enrichment in WT HIV-1). Residues were considered significantly remodeled when the 95% confidence interval for Δ(*WT — NAM*) excluded zero across the three independent simulation replicas. Changes in node weight, strength, and betweenness were analyzed to identify alterations in communication throughput, intensity, and relay, respectively.

**Figure 4.**
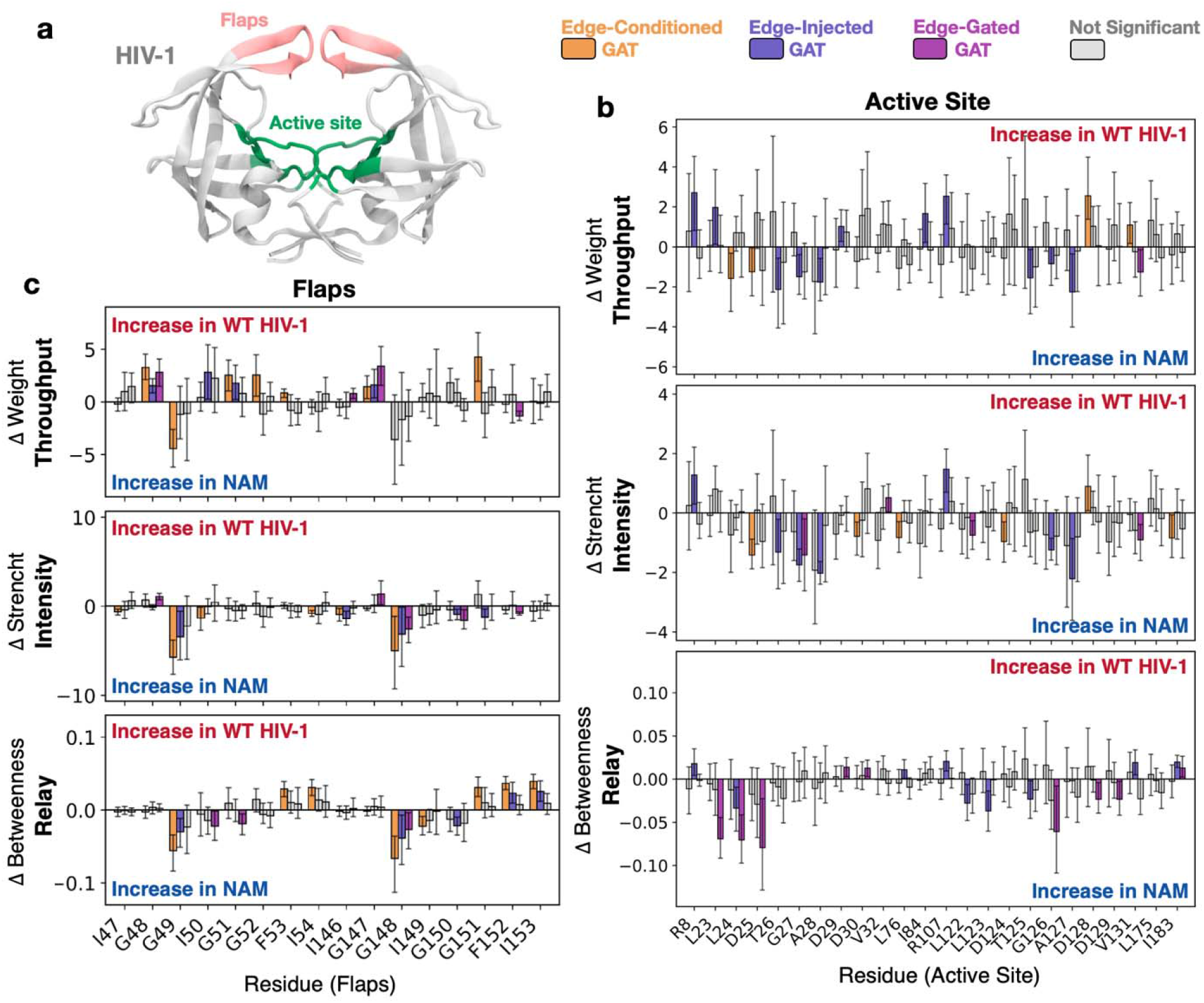
NAM-induced remodeling of residue communication in the flap and active site regions of HIV-1 protease. **(a)** Structure of HIV-1 protease highlighting the flap (pink) and active site (green) regions. **(b-c).** Residue-level differences in node weight (top), node strength (center), and node betweenness (bottom) between the wild-type (WT) HIV-1 and the non-active-site mutant (NAM) protease, , for **(b)** active site and **(c)** flap residues. The three communication descriptors – node weight, strength and betweenness – measure communication throughput, intensity and relay, respectively. Negative *Δ*(*WT* — *NAM*) indicate enrichment in NAM, whereas positive values indicate enrichment in WT HIV-1. Results are shown for the three edge-aware models: Edge-Conditioned, Edge-Injected, and Edge-Gated. Colored bars indicate statistically significant *Δ*(*WT* — *NAM*) for the corresponding model, whereas gray bars indicate residues that do not reach statistical significance. Bars represent the mean *Δ*(*WT* — *NAM*) across three independent simulations, and error bars denote the corresponding 95% confidence interval. Residues were considered significant when the confidence interval excluded zero.

Although none of the NAM substitutions occur within the active site, all three models detected significant communication remodeling at active-site residues, albeit with different descriptor-specific signatures (**Figure 4b**). The Edge-Conditioned model highlighted communication throughput and intensity in NAM, as indicated by negative Δ(*WT — NAM*) values for node weight and strength at residues including L24, D25, D30, and D124. The Edge-injected model produced a broader response in the catalytic region, with NAM enriched communication throughput and intensity across T26–A28 and T125–A127, together with communication relay (i.e., betweenness) at L24, L122, L123, and T125. The Edge-gated model primarily highlighted communication relay at the L23, L24, D25, G126, D128, and D129 residues. Collectively, these results show that the three edge-aware models detect allosteric communication remodeling within the catalytic region, despite the absence of active-site mutations, consistent with previous studies of HIV-1 protease allostery.^35–37^

The flap region showed a distinct descriptor pattern (**Figure 4c**). Across the three models, the flap residues showed little NAM enrichment in node weight (i.e., negative Δ(*WT — NAM*) values), indicating that the mutant does not establish the flap as a persistent communication throughput route. Instead, all three models showed NAM-enriched strength at selected flap residues, indicating increased communication intensity when those flap-associated edges are present. Representative NAM-enriched residues include G49/I50/I54/I146/G148 in the Edge-Conditioned model, and G148/I150/F152 in the Edge-Gated model. This pattern suggests that the NAM mutant does not establish the flaps as stable communication hubs but rather strengthens communication through specific flap-associated interactions when they form. This behavior is consistent with the increased flap flexibility and weakened regulation of flap dynamics previously reported in the NAM mutant.^20,33,34^

### Analysis of the mutation sites themselves revealed architecture-specific communication responses

The Edge-Conditioned model identified the largest number of significant mutation sites (12 of 18) across the three communication descriptors. The Edge-Injected model detected fewer mutation sites, with the clearest NAM-enriched signal at I15, which exhibited increased node weight and strength. The Edge-Gated model predominantly highlighted the I15/K20/M36 mutation cluster, consistent with its reported role in redistributing communication toward the flap hub in the multidrug-resistant NAM protease.^20^

Together, these residue-level analyses demonstrate that mutation-induced communication remodeling preferentially occurs within functionally important regions of HIV-1 protease, rather than being randomly (or uniformly) distributed across the structure. The Edge-conditioned model primarily detected communication throughput and intensity, highlighting local active-site/flap interactions as they form. The Edge-injected model captured broader redistribution of communication across functional and scaffold-associated residues. The Edge-Gated model most clearly resolved communication relay reorganization associated with allosteric propagation, linking the I15/K20/M36 mutation cluster to relay reorganization around the catalytic region.

### Region-level attention descriptors reveal architecture-specific NAM remodeling patterns

To determine whether NAM-induced communication remodeling extended beyond individual residues, we aggregated the residue-level communication descriptors into predefined structural regions and compared their values in WT HIV-1 and NAM. For each region, we calculated *b(wT — NAM)*, where negative values indicate increased communication in NAM and positive values indicate the opposite (**Figure 5**). HIV-1 is a symmetric homodimer, yet NMR and MD studies have shown that it frequently samples dynamically asymmetric conformations, including asynchronous flap motions^38–40^. Accordingly, the two symmetry-related monomers, conventionally referred to as the primed (′) and non-primed subunits, were analyzed separately to preserve potential dynamical asymmetries.

**Figure 5.**
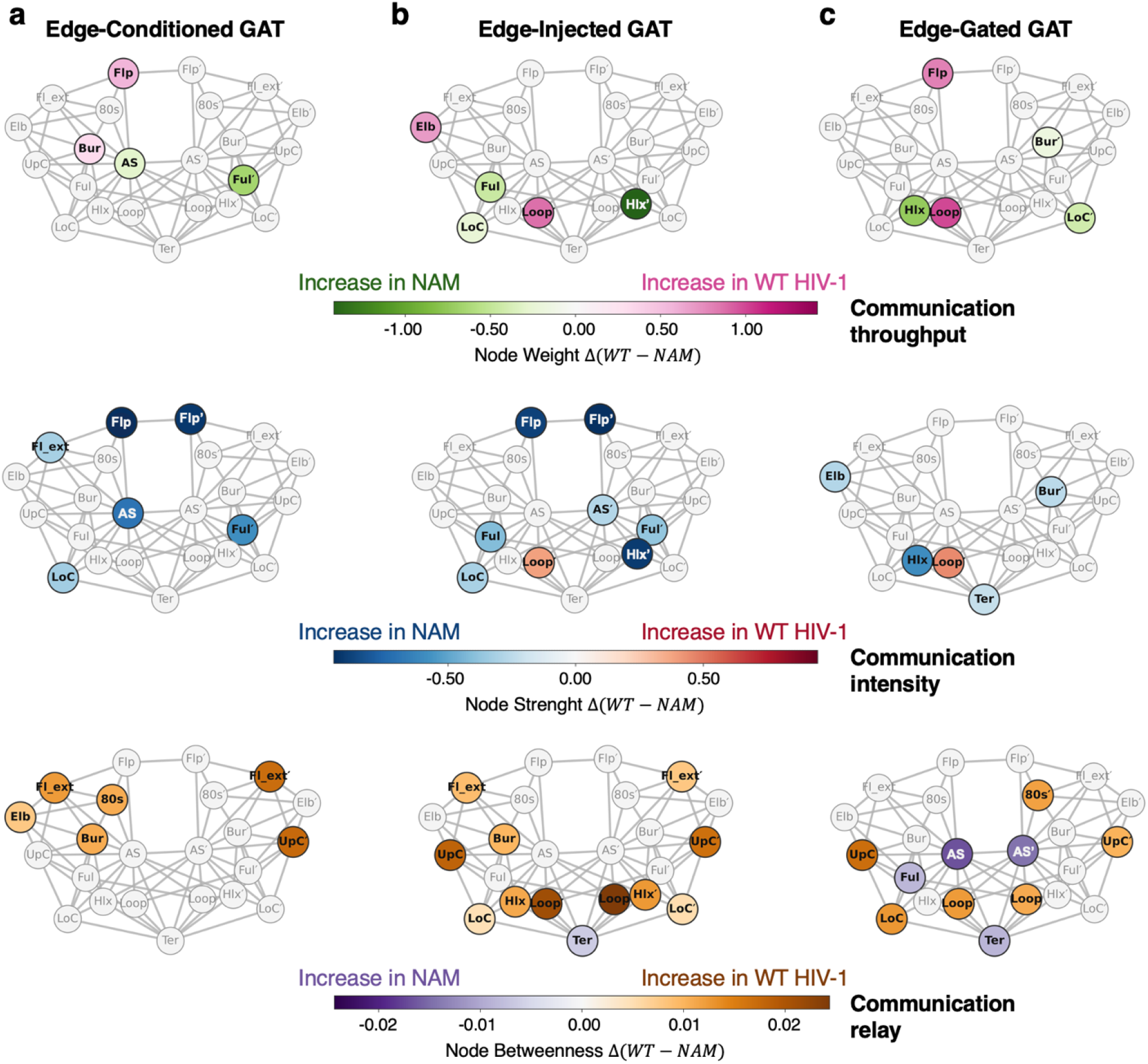
Region-level communication remodeling induced by non-active-site (NAM) mutations in HIV-1 protease. Communication networks comparing the wild-type (WT) HIV-1 and the NAM mutant for the three edge-aware models: **(a)** Edge-Conditioned, **(b)** Edge-Injected, and **(c)** Edge-Gated. Protein regions are represented as circles positioned according to a two-dimensional projection of their three-dimensional centers of mass calculated from the HIV-1 protease structure (PDB 1HHP)^41^. Gray edges denote structural contacts between regions (minimum interatomic distance ≤ 4.0 Å) shown as a fixed structural reference. Circles are colored according to significant *Δ*(*WT* — *NAM*) values for communication throughput (node weight, top), intensity (strength, middle), and relay (betweenness, bottom). Negative values indicate increased communication in NAM, whereas positive values indicate increased communication in WT. Colored nodes denote statistically significant *Δ*(*WT* — *NAM*), whereas gray nodes indicate non-significant regions (95% confidence interval includes zero).

The Edge-Conditioned model revealed NAM-induced remodeling centered on the functional regions of HIV-1 protease, namely the active site and flap regions (**Figure 5a**). The active site (AS) and fulcrum (Ful′) exhibited increased node weight and strength in NAM (i.e., Δ < 0, top and central panels), indicating enhanced communication throughput and intensity. The flap regions (Flp/Flp′) displayed reduced communication throughput but increased intensity, indicating that although the flaps participate less consistently in the communication network, the interactions they establish become more communication-intensive when they occur.

This pattern is consistent with the increased flap flexibility and disrupted flap communication previously reported for NAM variants.^20,33,34^ In contrast, communication relay (node betweenness, bottom panel) generally decreased in NAM, with no redistribution at the active site and flap regions. Thus, the Edge-Conditioned model primarily highlighted mutation-induced remodeling of the functional regions through changes in communication throughput and intensity rather than communication relay.

The Edge-Injected model GAT_EIMP_ captured a distinct pattern of NAM mutation-induced communication remodeling (**Figure 5b**). The functional regions, including both flaps (Flp/Flp’) and active site (AS), were primarily enriched in node strength (central panel), indicating increased communication intensity rather than sustained communication throughput across the trajectory. In contrast, scaffold-associated regions, including the fulcrum (Ful), lower cantilever (LoC), and helix (Hlx′), exhibited increased node weight and strength in NAM (top and central panels, respectively), indicating communication throughput and intensity across the structural scaffold. Node betweenness (bottom panel) further suggested broad attenuation of the communication relay in NAM. Together, these observations indicate that the Edge-Injected model primarily captures communication intensity within the functional regions while distinguishing it from the sustained communication throughput maintained by the structural scaffold. This pattern is consistent with the increased flap flexibility previously reported for NAM variants in computational and biophysical studies.^20,40^

The Edge-Gated model primarily highlighted NAM-induced remodeling of communication relay (**Figure 5c**). Unlike the other models, the catalytic and flap regions in NAM were not broadly enriched in both communication throughput and intensity. Instead, increased node weight and strength were primarily observed within scaffold-associated regions, particularly the helix (Hlx) and buried (Bur′) regions. Most notably, node betweenness (bottom panel) redistributed toward the active site (AS), fulcrum (Ful), and terminal (Ter) regions, indicating pronounced reorganization of the communication relay. These observations indicate that the Edge-Gated model primarily captures mutation-induced remodeling through communication relay, highlighting long-range relay pathways centered on the functional regions of the protease.

## Discussion

Deep learning is enabling increasingly accurate prediction of biomolecular structures, molecular properties, energies, and forces, and is emerging as a powerful tool for analyzing MD simulations.^13–16,42,43^ While these models are usually evaluated by predictive accuracy, the organization of the learned dynamics remains difficult to interrogate mechanistically.

Here, we introduce three edge-aware GAT models that reproduce coordinate evolution within sampled MD trajectories with sub-angstrom accuracy and enable mechanistic interpretation of biomolecular dynamics through attention-derived communication networks. The methodological contribution of this work is the integration of coordinate prediction, perturbation-based faithfulness analysis, and attention-derived network analysis within a unified approach. We introduce three edge-aware message passing models – Edge-Conditioned, Edge-Injected and Edge-Gated – that differ in how edge features contribute to message passing: by biasing attention coefficients, modifying transmitted messages, or gating information transfer (**Figure 1c**). This controlled design enabled us to isolate the effect of edge-aware message passing on both predictive performance and mechanistic interpretation. Rather than interpreting attention as an explanation by itself,^17,18^ we first established through perturbation-based faithfulness analysis that highly attended edges contributed more strongly to prediction than randomly selected edges. We then characterized the resulting communication networks using three complementary communication descriptors – node weight, strength and betweenness – which measure communication throughput, intensity, and relay, respectively.

Chignolin provided a simple benchmark for mechanistic interpretation because its β-hairpin folding is governed by well-defined structural determinants (**Figure 2a**). The Tyr2-Trp9 aromatic interaction and the Asp3/Gly7/Thr8 turn-associated network are known to stabilize the native fold.^19,31,32^ The three edge-aware models report comparable coordinate prediction accuracy (**Figure 2b,c**) and recovered distinct communication patterns (**Figure 2e-g**). The Edge-Conditioned model emphasized the Trp2 aromatic residue as a dominant communication hub. The Edge-Injected model separated communication hubs from relay residues, with the Tyr2-Trp9 aromatic interaction dominating communication throughput and intensity, whereas the Gly7/Asp3/Thr8 turn region was preferentially identified as the communication relay. The Edge-Gated model most clearly resolved the Asp3/Gly7/Thr8 turn-stabilizing network while preserving strong communication through the Tyr2-Trp9 interaction. Collectively, these findings demonstrate that different formulations of edge-aware message passing recover complementary mechanistic aspects of residue communication.

HIV-1 protease provided a more complex test case because its function depends on long-range communication, rather than a small number of local structural determinants.^19,34,40,44,45^ Unlike Chignolin, communication is distributed across multiple functional regions and is extensively remodeled by non-active-site resistance mutations (NAM, **Figure 3a**).^35–37^ Previous studies have shown that these mutations remodel the functional regions of HIV-1 protease by altering active-site communication, increasing flap flexibility, weakening regulated flap communication, and redistributing communication through scaffold-associated pathways.^20^ These established allosteric signatures therefore provide a stringent benchmark for assessing whether the communication networks recovered by the three edge-aware models capture biologically meaningful remodeling of allosteric communication.

The three edge-aware models captured complementary aspects of this established allosteric mechanism, consistent with their different message-passing designs (**Figures 4,5**) The Edge-Conditioned model highlighted communication remodeling within the functional regions (i.e., the catalytic site and flaps) through changes in communication throughput and intensity rather than communication relay (**Figure 5a**). The Edge-Injected model identified NAM-induced remodeling in which the catalytic and flap regions were primarily characterized by increased communication intensity rather than sustained communication throughput (**Figure 5a**), while communication throughput and intensity were redistributed toward scaffold-associated regions. Compared to WT HIV-1, broad attenuation of communication relay was observed, rather than the emergence of a dominant relay pathway in the mutant. The Edge-Gated model most clearly resolved communication relay within the functional active-site region following NAM remodeling (**Figure 5c**). Indeed, while communication throughput and intensity largely remained confined to scaffold-associated regions, relay organization was redistributed toward the catalytic region, fulcrum, and terminal interface.

Together, these findings demonstrate that the three edge-aware models provide complementary views of residue communication rather than a single, unique mechanistic interpretation. Because the three models differed in how edge features contributed to message passing, the complementary communication networks recovered across Chignolin and HIV-1 protease show that the mechanistic interpretation extracted from graph attention depends on how edge information is incorporated into message passing (**Figure 6a**). The Edge-Conditioned model preferentially emphasizes functional regions characterized by high communication throughput and intensity. The Edge-Injected model highlights regions characterized by high communication intensity, such as the flap regions in HIV-1 protease and key stabilizing interactions in Chignolin, identifying interactions that contribute strongly to communication when present. The Edge-Gated model most clearly resolves long-range communication relay ions, identifying residues that serve as bridges or bottlenecks linking distant communication pathways.

**Figure 6.**
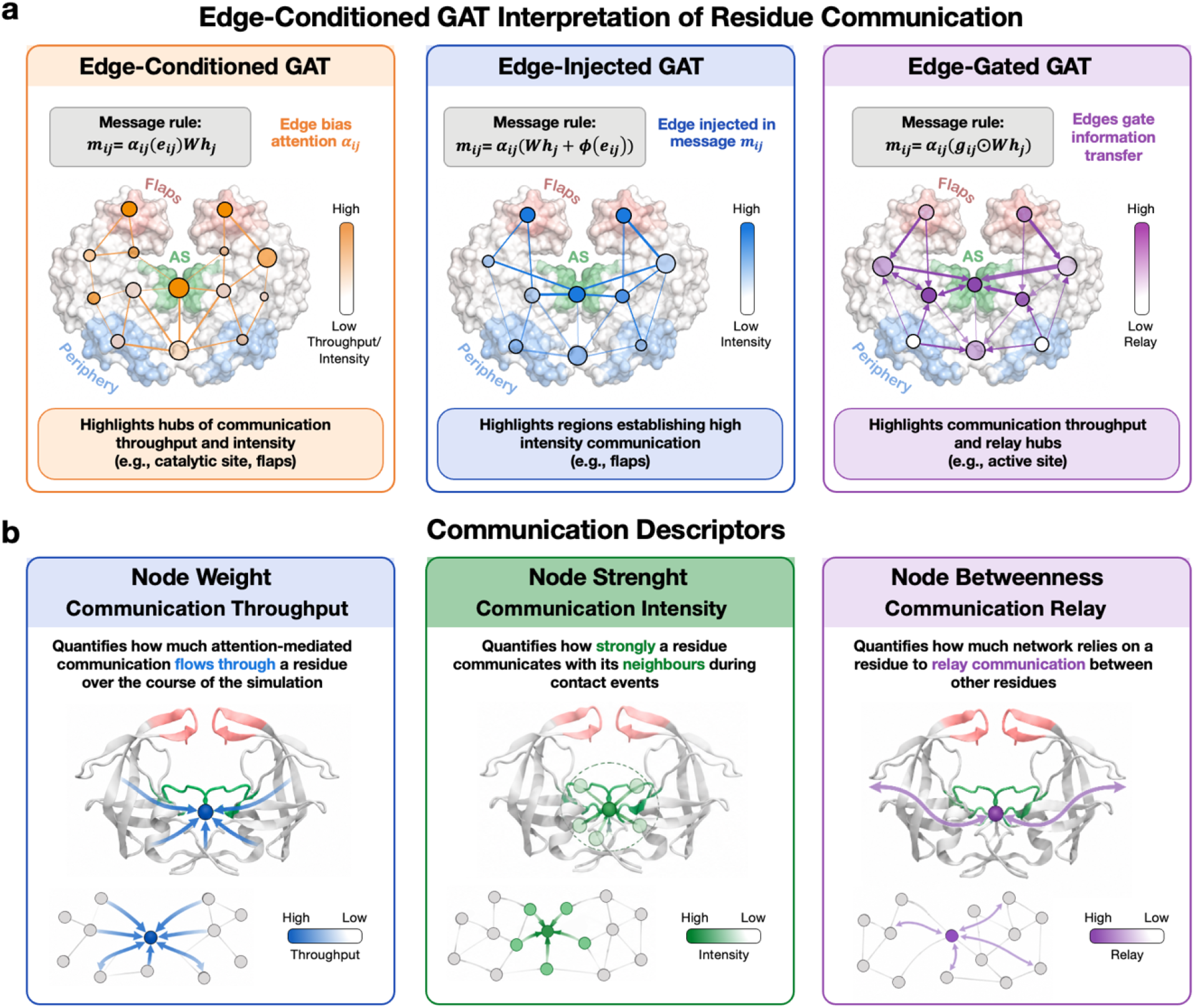
Mechanistic interpretation of residue communication using edge-aware graph attention networks (GAT). **(a)** The three edge-aware GATs – Edge-conditioned (left), Edge-Injected (center), and Edge-Gated (right) – differ in how edge information is incorporated during message passing, yielding complementary interpretations of residue communication. The Edge-Conditioned GAT identifies communication hubs characterized by high communication throughput and intensity. The Edge-Injected GAT highlights regions establishing high-intensity communication, identifying interactions that contribute strongly to communication when present. The Edge-Gated GAT most clearly resolves long-range communication relay, identifying residues that mediate communication between distant regions of the network. **(b)** Attention-derived communication networks are characterized using three complementary graph descriptors. Node weight (left), node strength (center), and node betweenness (right) quantify communication throughput, intensity, and relay, respectively, enabling characterization of complementary communication roles within biomolecular networks. Together, the edge-aware GAT models and communication descriptors provide an interpretable AI framework for identifying functional regions that sustain communication throughput, establish high-intensity communication, or mediate long-range communication relay from molecular dynamics simulations.

To extract mechanistic information from these models, we quantified the learned communication networks using three descriptors – node weight, strength, and betweenness – describing communication throughput, intensity, and relay, respectively (**Figure 6b**). Together, the three edge-aware models and the corresponding communication descriptors provide a comprehensive framework for identifying functional regions characterized by persistent communication throughput, high communication intensity, and long-range communication relay.

The present approach also identifies several opportunities for future development. First, the current models were developed and evaluated for one-step prediction within sampled MD trajectories. Their applicability to autonomous trajectory generation, as well as to proteins and mutation classes not represented in the training set, remains to be established. Second, the current graph representation is intentionally simple, using Cα residues as nodes and distance-based edge features, without explicitly representing directional interactions, hydrogen bonding, electrostatics, solvent effects, or residue-specific physicochemical properties. Incorporating richer structural and physicochemical edge representations represents a natural next step toward improving both predictive accuracy and mechanistic interpretability. More broadly, extending this approach to multi-step trajectory prediction, transfer across diverse biomolecular systems, and richer molecular graph representations will further expand the scope of interpretable graph attention models for molecular dynamics simulations.

Overall, the edge-aware GAT models presented here provide complementary interpretations of biomolecular dynamics, with each model emphasizing distinct aspects of residue communication. This approach provides a general strategy for characterizing the communication networks that underlie biomolecular function, including folding, allostery, and long-range signal propagation in molecular machines. Allosteric communication is now recognized as a fundamental property of biomolecules, governing function, specificity, and regulation across a wide range of enzymes and molecular machines^46–52^. Our previous computational studies of gene editing systems illustrate the importance of identifying residues that mediate long-range communication. In CRISPR–Cas9, characterization of the allosteric communication network identified key residues responsible for transmitting the DNA-binding signal to the catalytic sites,^53^ and experimental perturbation of these communication hubs led to variants with improved genome-editing specificity^54–56^. Similarly, in the RNA-targeting nuclease Cas13a, analysis of the allosteric communication network identified residues controlling long-range signaling^57^, whose engineering yielded variants with enhanced RNA-targeting specificity for molecular detection and imaging applications^58^.

These examples illustrate that identifying residues central to allosteric communication is important not only for understanding biomolecular function but also for guiding protein engineering. Beyond identifying communication hubs, however, distinguishing whether a residue primarily sustains communication throughput, mediates local communication intensity, or serves as a communication relay provides a richer mechanistic basis for prioritizing mutational targets. By identifying communication hubs, relay pathways, and regions of sustained or transient communication, the edge-aware GAT framework introduced here offers a systematic and interpretable strategy for AI-guided identification of residues for functional perturbation and rational protein engineering.

## Conclusions

Here, we introduced an interpretable AI approach that combines molecular dynamics prediction with mechanistic interpretation through attention-derived communication networks. By explicitly incorporating edge information into message passing, our approach improved coordinate prediction relative to a standard graph attention network (GAT), demonstrating the importance of edge-aware learning for biomolecular dynamics. At the same time, our three edge-aware models – Edge-Conditioned, Edge-Injected and Edge-Gated – enabled mechanistic interpretation of learned molecular representations, recovering complementary aspects of residue communication. The Edge-Conditioned model preferentially identified communication hubs characterized by high throughput and intensity, the Edge-Injected model highlighted functional regions with high communication intensity, and the Edge-Gated model most clearly resolved long-range communication relay through functionally important regions. Together with graph-theoretical descriptors of the learned communication networks – node weight, strength, and betweenness – these complementary views provide a systematic approach for characterizing communication throughput, intensity, and relay from MD simulations. More broadly, this work demonstrates how interpretable AI can move beyond trajectory prediction toward mechanistic understanding of biomolecular dynamics. By providing complementary views of residue communication, this edge-aware GAT approach offers a general strategy for investigating folding, allostery, and long-range signaling in biomolecular systems, while enabling AI-guided identification of communication hubs and relay pathways for functional perturbation and rational protein engineering.

## Acknowledgments

This material is based upon work supported by the NIH (Grant No. R35GM164142 to G.P.) and the NSF (Grant No. CHE-2144823 to G.P.). GP acknowledges support by the Camille and Henry Dreyfus Foundation (Grant No. TC-24-063). The computational studies performed here were carried out using Bridges2 at the Pittsburgh Supercomputer Center through allocation BIO230007 from the Advanced Cyberinfrastructure Coordination Ecosystem: Services & Support (ACCESS) program, which is supported by NSF support grants #2138259, #2138286, #2138307, #2137603, and #2138296.

## biorxivCompeting Interests

The authors declare no competing financial interests.

